# Caerin 1.1 and 1.9 peptides from Australian tree frog inhibit antibiotic-resistant bacteria growth in a murine skin infection model

**DOI:** 10.1101/2021.04.20.440726

**Authors:** Shu Chen, Pingping Zhang, Liyin Xiao, Ying Liu, Kuihai Wu, Guoying Ni, Hejie Li, Tianfang Wang, Xiaolian Wu, Guoqiang Chen, Xiaosong Liu

**Affiliations:** Cancer Research Institute, Foshan First People’s Hospital, Foshan, Guangdong 528000, China; Clinical Microbiological Laboratory, Foshan First People’s Hospital, Foshan, Guangdong 528000, China; Genecology Research Centre, University of the Sunshine Coast, Maroochydore DC, QLD 4558, Australia; Department of Rheumatology, Foshan Frist People’s Hospital, Foshan, Guangdong 528000, China

**Author notes:** **Corresponding to:** Dr. Guoqiang Chen, Dr. Xiaosong Liu. Shu Chen and Pingping Zhang contributed equally to this work. Author order was determined in the alphabetical order of their last names.

## Abstract

Host-defence caerin 1.9 peptide was originally isolated from skin secretion of Australian tree frog, and inhibits the growth of a wide range of bacteria *in vitro*. In this study, we demonstrated that caerin 1.9 shows high bioactivity against several bacteria strains, such as *Staphylococcus aureus, Acinetobacter Baumannii*, methicillin-resistant *Staphylococcus aureus* (MRSA), and *Streptococcus hemolyiicus in vitro*. Importantly, unlike antibiotic Tazocin, caerin 1.9 does not induce bacterial resistance after 30 rounds of *in vitro* culture. Moreover, caerin 1.1, another peptide of caerin family, has additive antibacterial effect when used together with caerin 1.9. Furthermore, caerin 1.1 and 1.9 prepared in the form of a temperature sensitive gel inhibit *MRSA* growth in skin bacterial infection model of two murine strains. These results indicate that caerin 1.1 and 1.9 may have the advantage than conventional antibiotics against bacterial infection of skin.

## Introduction

Skin and soft tissue infections (SSTIs) are resulted from pathogenic invasion of the skin and underlying soft tissues and have variable presentations, aetiologies and severities (1). The estimated incidence rate of SSTIs is 24.6 per 1000 person-years (2), and approximately 70% to 75% of all cases are managed in the outpatient setting (3). *Staphylococcus aureus* is the leading cause of both uncomplicated and complicated infections. Moreover, multidrug-resistant bacteria, mainly methicillin-resistant *S. aureus* (both community-acquired and healthcare-associated), are associated with significantly increased morbidity, mortality, length of hospital stay, and costs (4). SSTIs are generally managed in the community, including self-apply of antibiotics. Oral or systemic use of antibiotics is recommended at outpatients or in hospital, depending on the severity of the symptoms. Currently, topical application of antibiotics, such as mupirocin is recommended for mild impetigo and folliculitis (5).

Due to the misuse and overuse of antibiotics, the emergence of antibiotic-resistant bacteria is becoming a serious health challenge (6). The rate of antimicrobial-resistant infections grows dramatically. Therefore, a novel and effective antibacterial agent is essential in the nowaday community.

Naturally derived host defense peptides are one of the first successful forms of defense of eukaryotes against bacteria, protozoa, fungi, and viruses. More than 200 host-defense peptides have been isolated and identified from skin secretions of Australian frogs and toads. Many of these peptides, including caerin peptides, have antiviral, antitumor, antimicrobial, and/or neuropeptide-type activities (7-10). Caerin 1.1 has an anti-cancer effect against a number of human cancer cell lines, including leukaemia, lung, colon, CNS, melanoma, ovarian, renal, prostate, and breast cancers (8, 11). Substitution of either or both of the Pro residues with Gly leads to the peptide with overall reduced activity (12). Both caerin 1.1 and caerin 1.9 peptide has antimicrobial activity against a wide spectrum of Gram-positive and Gram-negative microbial strains *in vitro* (7, 13-15). Caerin 1.1 and 1.9 inhibit HIV-infected T cells within minutes post-exposure at concentrations non-toxic to T cells and inhibit the transfer of HIV from dendritic cells (DCs) to T cells with limited toxicity (10, 16). Recently, it has been shown that Caerin 1.1 and Caerin 1.9 peptides have additive effects against human papillomavirus transformed tumour cells and increase the efficacy of a therapeutic vaccine against HPV related diseases (9, 17-19).

In this study, we investigated whether caerin 1.9 peptide is able to inhibit antibiotic-resistant bacteria growth and whether caerin 1.1 and caerin 1.9 have additive effect against bacteria growth *in vitro*. Their anti-bacterial ability was examined in a skin bacterial infection model *in vivo*.

## Results

### Minimum Inhibitory Concentration (MIC) of standard bacterial strains by caerin 1.9

We first performed the MICs of caerin 1.9 against *S. aureus, P. aeruginosa*, MRSA, *A. Baumannii*, and *S. hemolyiicus*. Bacteria in the logarithmic phase were cultured in the presence of different concentration of caerin 1.9. The final concentrations of caerin 1.9 were 120, 60, 30, 15, 7.5, 3.75, and 1.875µg/ml respectively. The MIC was defined as the concentration of the caerin 1.9 in which bacterial growth is completely inhibited. The MICs of *S. aureus*, MRSA, *A. Baumannii*, and *S. hemolyiicus. Bacteria* were 7.5µg/ml, while the MIC of P. aeruginosa was 60µg/ml by caerin 1.9 (**Table 1** and **Table S)**.

**Table 1.**
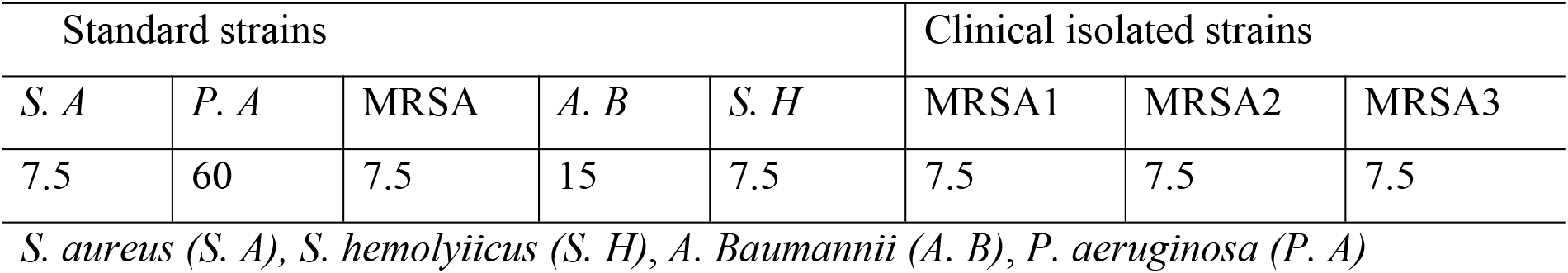
Minimum inhibitory concentration MIC (µg/ml) of caerin1.9 against standard and clinically isolated bacteria strains.

We next investigated the inhibitory effect of caerin 1.9 on clinically isolated MRSA. The patients’ information and antibiotic resistance of these bacteria strains were shown in **Table S2**. The MICs of caerin 1.9 against clinically isolated MRSA samples remain at 7.5 μg/ml (**Table 1**).

To better understand the dynamic inhibitory actions of caerin 1.9 against the growth of *S. aureus, P. aeruginosa*, MRSA, *A. Baumannii*, and *S. hemolyiicus*, these bacteria were then exposed to caerin 1.9 with the concentration of MIC or 1/4 MIC. When the caerin 1.9 was co-cultured with the bacteria at the MIC value, the bacterial growth was completely inhibited in all tested bacteria for 24 hours, except for *P. aeruginosa*. When the bacteria were co-cultured with 1/4 of the MIC, the bacteria began to grow after 6 hours. While the bacteria in the PBS group started to grow after 4 hours. Meanwhile, the growth of *S. hemolyiicus* was slower than other bacteria, which started increasing after 10-hour culture for both PBS and 1/4MIC group. After 48-hour cultivation with MIC, caerin 1.9 still inhibited *S. aureus*, MRSA, *A. Baumannii*, and *S. hemolyiicus*, but the inhibition of *P. aeruginosa* was poor (**Figure 1**).

**Figure 1.**
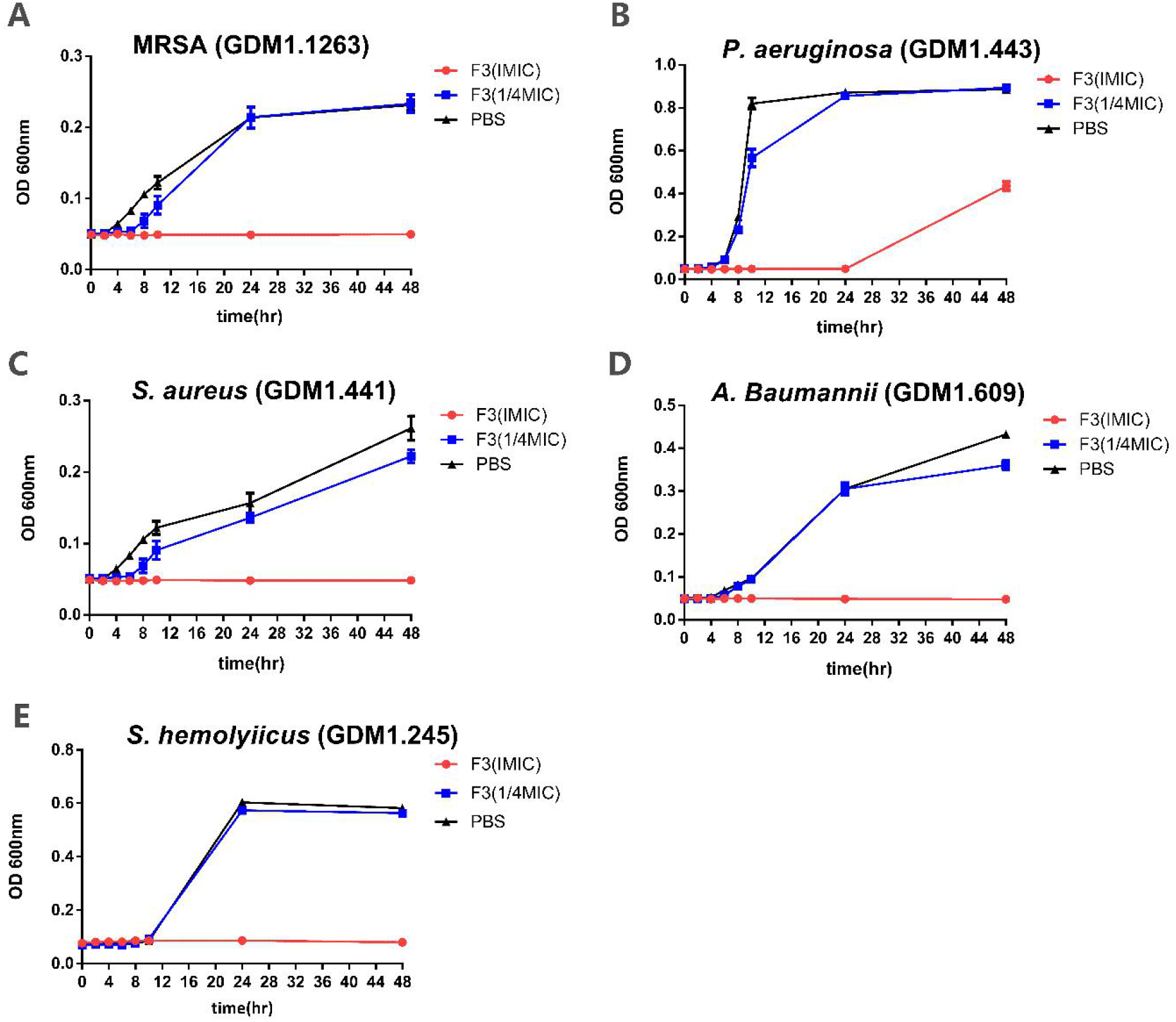
Change of optical density for *S. aureus, P. aeruginosa*, MRSA, *A. Baumannii*, and *S. hemolyiicus*. Different Bacteria at the OD of 0.08-0.1were cultured in media with PBS or with caerin 1.9 at MIC or 1/4 MIC. OD values were measured at 0, 2, 4, 6, 8, 10, 24, 48hr by UV spectrometer at the wavelength of 600nm.

### Caerin1.9 dose not induce resistance strains

Next, we investigated whether Caerin 1.9 induces bioresistant strains. The logarithmic *P. aeruginosa* and MRSA were cultured in 96-well plates containing caerin1.9 or piperacillin sodium (Tazocin sodium) at one-fourth MIC. The medium was replaced every three days, and the whole culture process was continued for three months. After three months of culture with 30 passages, the MIC of caerin1.9 against MRSA and *P. aeruginosa* did not change, suggesting no drug-resistant strain was induced. While the MIC of the antibiotic Tazocin incrased by a16-fold in MRSA and an 8-fold for *P. aeruginosa* (**Table 2**).

**Table 2.**
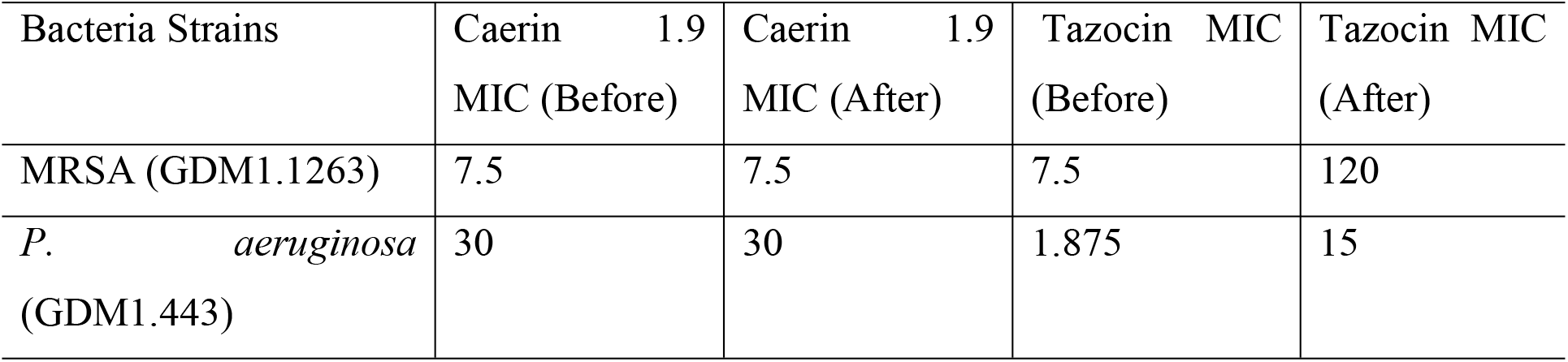
Comparison of caerin1.9 and Tazocin on induced resistance of MRSA and *Pseudomonas aeruginosa*.

### Inhibition of S. aureus, MRSA and S. hemolyiicus by caerin 1.1/caerin 1.9 using disc diffusion method

As *S. aureus*, MRSA and *S. heolyiicus* are most oftenly isolated bacteria strains during skin infection, the anti-bacterial ability of caerin 1.1 and 1.9 against bacterial strains that identified mostly from skin infection were tested by using the disc diffusion method. Standard *S*.*aureus*, MRSA, and *S*.*hemolyiicus* strains were used. Polymyxin B discs were used as a control, as it was an antibiotic used against skin infection, which is capable to induce a visible inhibiton zone in all media at the dose of 120µg or above **(Table 3)**.

**Table 3.**
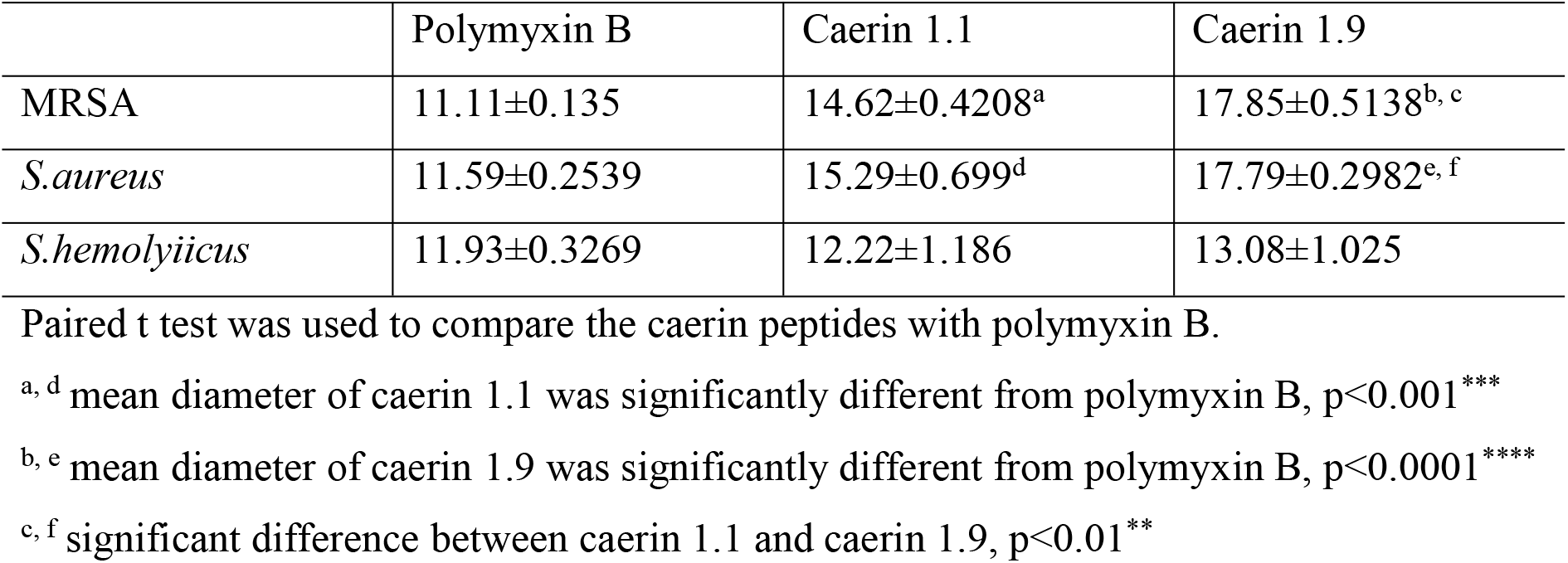
Comparison of inhibition zone diameters among polymyxin B, caerin 1.1, and caerin 1.9 at a dose of 120µg

The inhibition diameters of polymyxin B were consistent among three types of bacteria, at 11-12mm. Caerin 1.1 or caerin 1.9 developed larger inhibition zones, up to around 15mm or 18mm against MRSA as well as *S. aureus*. For the inhibition zones developed in the *S. hemolyiicus* media, no significant difference between caerin 1.1 and polymyxin B or between caerin 1.9 and polymyxin B. The results suggested that both caerin 1.1 and caerin 1.9 have a better ability to inhibit the growth of *S. aureus* and MRSA than a commonly used polymyxin B, but not significant against *S. hemolyiicus*. Generally, caerin 1.9 exhibited a stronger antimicrobial activity compared to caerin 1.1.

However, due to the hydrophilicity of caerin peptides, they have poor abilities to diffuse in the agar plate (20). Thus, the disc diffusion method may not reflect their actual ability in bacterial inhibition. Also, the disc diffusion method only provided a qualitative test for these peptides, which was not able to demonstrate the relationship between peptide concentrations and zone diameters.

### Caerin 1.1 and caerin1.9 has additive effects against MRSA

The combined bacteria growth inhibition effect of caerin1.1 and caerin1.9 on MRSA and *A. Baumannii* was tested using the microdilution checkboard method (21). This method was based on the M38-A2 CLSI protocol for evaluating the bactericidal activity of the combination of caerin 1.1 and caerin 1.9 in different concentrations at 24hr. The in vitro interactions were calculated and interpreted as FICI. The FICI against both bacteria was 0.5 <FICI ≤ 1, suggesting that they have an additive antibacterial effect on MRSA and *A. Baumannii* (**Table 4**).

**Table 4.**
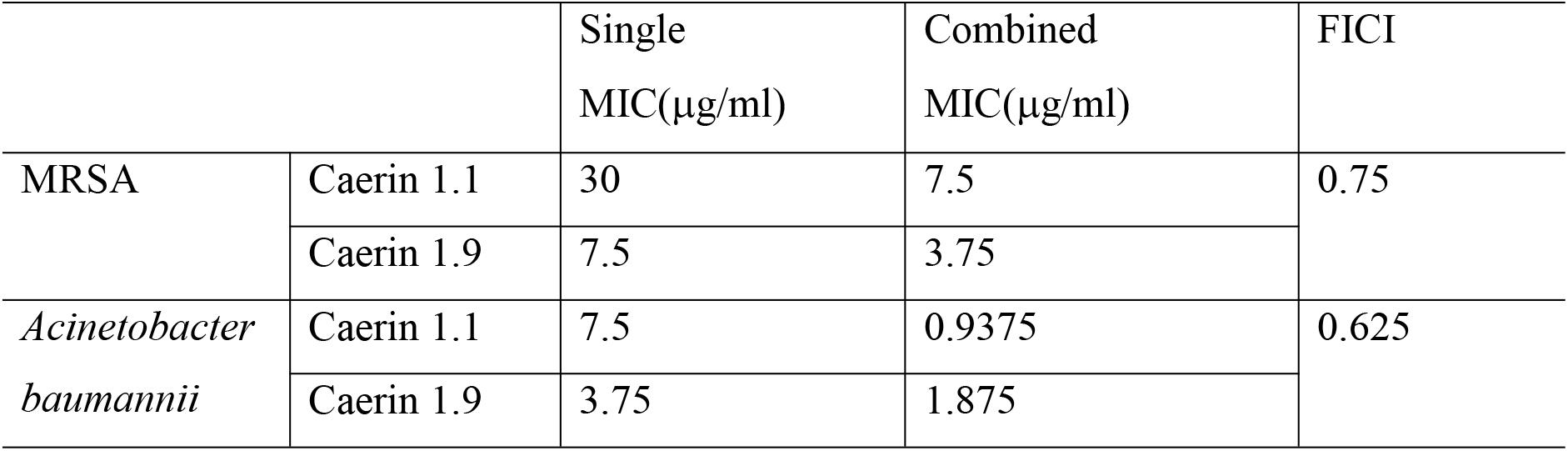
Caerin1.1 and caerin1.9 combination against MRSA and *Acinetobacter baumannii*.

### Caerin 1.1 and 1.9 inhibit skin MRSA growth in mice

Next, we investigated whether caerin 1.1 and caerin 1.9 inhibit bacteria growth in a skin MRSA infection model. We first tested whether the combinations of caerin 1.1 and caerin 1.9 (in an one-to-one molar ratio) inhibit the growth of MRSA isolated from mouse skin infection site. Pus materials resulted from MRSA skin infection was mixed with different concentrations of caerin gel, followed by inoculating onto a nutrition agar plate (LS0309, Guangzhou Dijing Microbiology Technology Co., Ltd.). After overnight incubation, the number of bacterial colonies indicated the *in vitro* anti-bacterial ability of caerin gel. At 1.5625mg/ml, caerin gel started to inhibit the growth of MRSA, and the caerin gel almost eradicated the present of MRSA at the concentration of 12.5mg/ml (**Figure 2**). Then, a severe skin damage model was established to ascertain that caerin peptide is able to achieve anti-bacterial activity *in vivo* (**Figure 3**).

**Figure 2.**
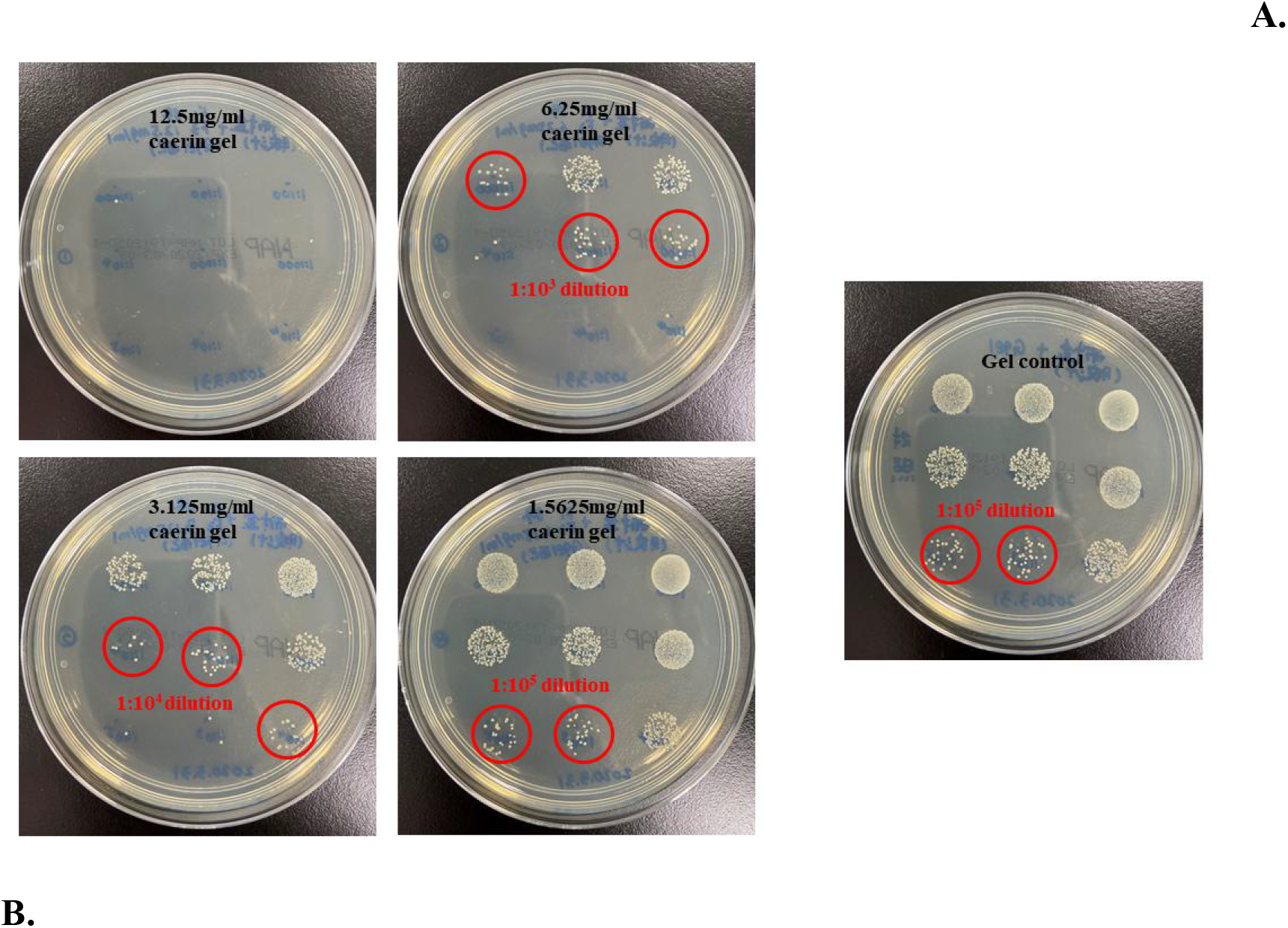
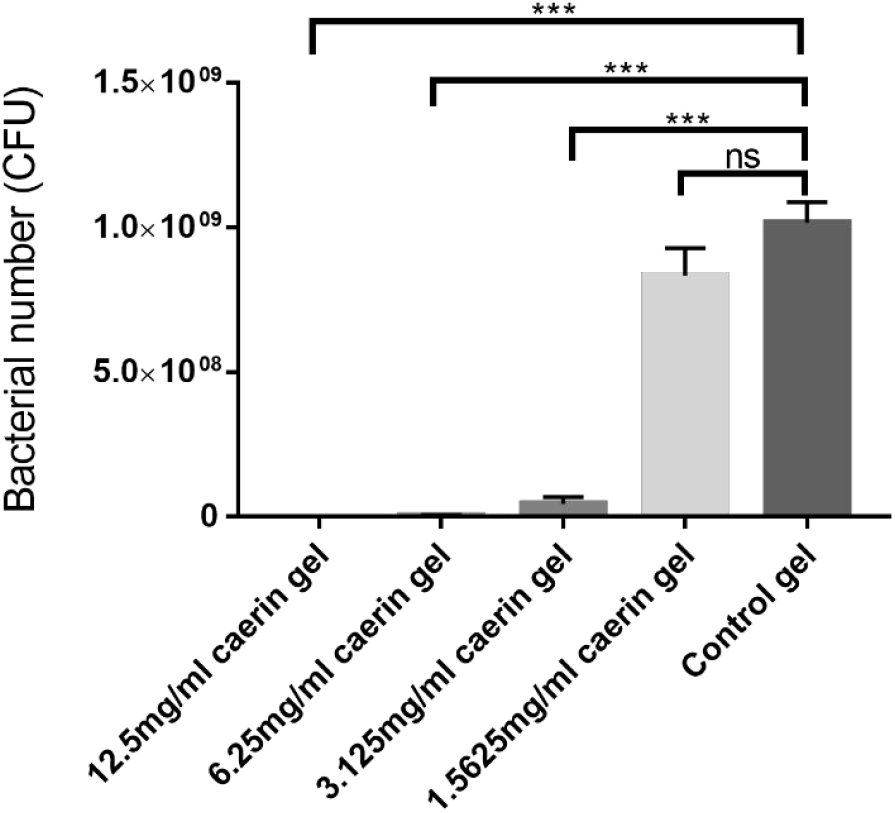
Caerin 1.1 and 1.9 gel inhibits the growth of MRSA from skin infection area *in vitro*. Pus resulted from MRSA infected murine skin was mixed with different concentrations of caerin gel (12.5 6.25, 3.125, and 1.5625 mg/ml) and control poloxamer gel, then 30ul of the mixture was put onto a nutrition agar plate and incubated at 37°C for 24 hours. Each bar represented the number of countable bacterial colonies at certain concentration, and the error bar represented the standard deviations. ***P<0.001; ns, not significant.

**Figure 3.**
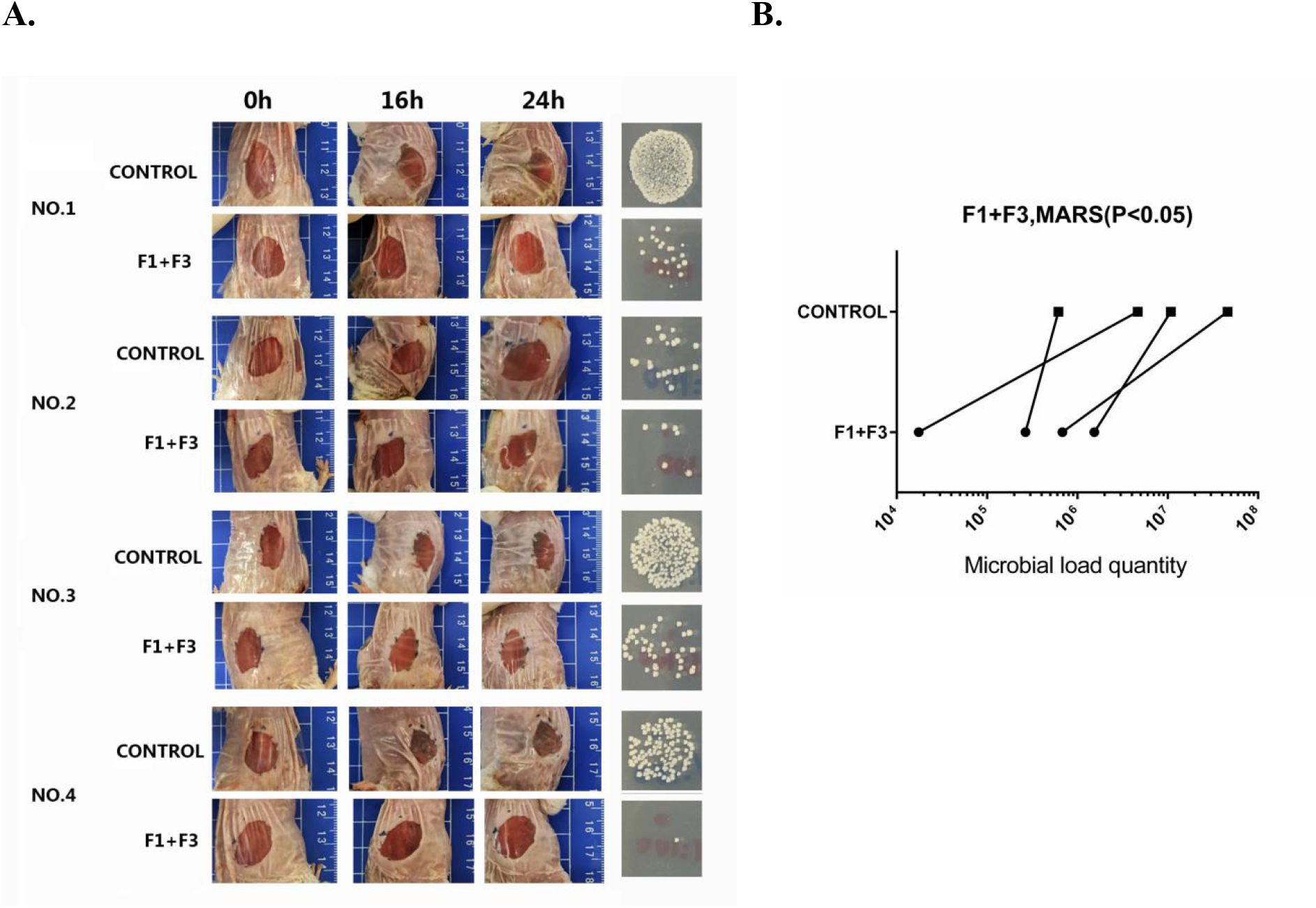
Balb/c mice with severe MRSA skin infection, and the pair comparisons of bacteria counts between caerin gel and control gel. Approximately 5×10^6^ CFU were inocultured on each wound; with 12.5mg/ml of caerin 1.1 (F1) and caerin 1.9 (F3) peptides in poloxamer gel, or with gel only for three days. Four mice were included in this group. The caerin gel inhibited the growth of MRSA at a significant level of p<0.05.

The tape-stripped infected model was developed for investigating the therapeutic effect of caerin gel on MRSA infected murine skin. The epidermal layer was damaged, allowing bacteria to colonize and grow on the skin surface as described in the Materials and Methods (22). Four hours after infection, 20µl of caerin gel or P3 gel was applied to one infection area, leaving another side with gel only for 2 times/day, 3 days. And a group in which both damaged sides were applied with saline water was designed as control. As a result, similar phenomena were observed in both Balb/c mice (**Figure 4**), C57BL/6 mice (**Figure 5**), and the severe MRSA skin infection model (**Figure 3**). The saline water provided no therapeutic effect for the MRSA infection (**Figure 4D, 5C**). The bacterial counts of the caerin gel treated areas were significantly reduced, compared to the untreated areas (P<0.05) (**Figure 4A, 5A**). In the P3 gel group, the P3 treated and untreated areas had a similar number of bacteria (**Figure 4C, 5B**). These results suggested that the caerin peptide gel was effective in inhibiting bacteria growth for the superficial skin infection caused by MRSA.

**Figure 4.**
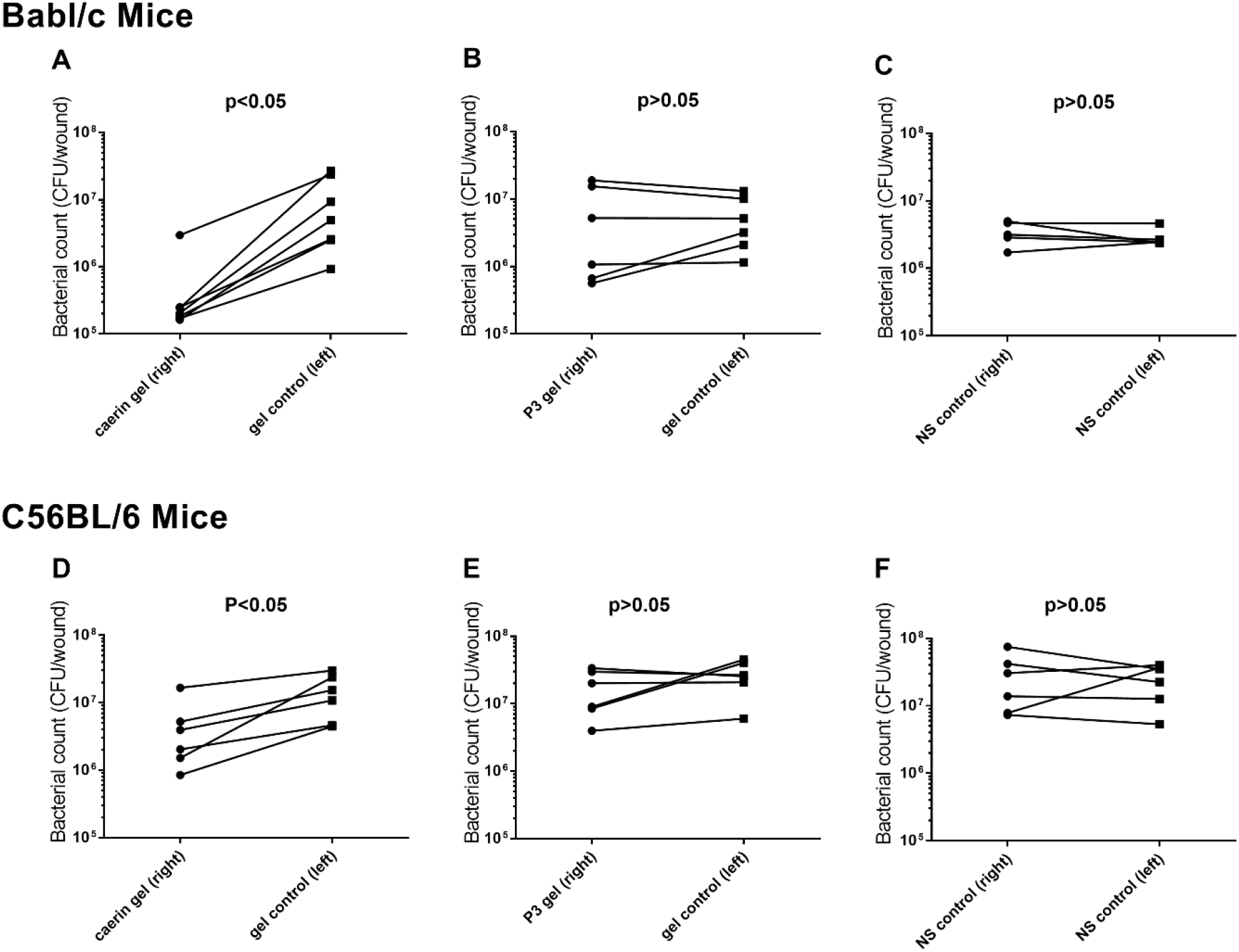
Caerin 1.1 and 1.9 gel inhibited the growth of MRSA (GDM1.1263). Tape-stripped Babl/c mice and C57BL/6 mice were infected with MRSA. Comparisons of bacteria counts between caerin gel (combined caerin 1.1 and caerin 1.9) and control gel, between P3 gel and control gel, and between two sides in the normal saline control group. Approximately 5×10^4^ CFU were inocultured on each wound; with 12.5mg/ml of caerin 1.1 and caerin 1.9 peptides in poloxamer gel, or with gel only for three days. Each line represented the pair comparison of the number of bacteria (CFU) between the right and left stripped areas per mice. **A**. The pair comparison of the CFU between caerin gel and control gel. **B**. The pair comparison of the CFU between P3 gel and control gel. **C**. The pair comparison of the CFU between two sides of normal saline. **D**. The pair comparison of the CFU between caerin gel and control gel. **E**. The pair comparison of the CFU between P3 gel and control gel. **F**. The pair comparison of the CFU between two sides of normal saline. The number of mice in each group was as followed: A, n=7; B, n=6; C, n=5; D, n=6; E, n=6; F, n=6. The result is the representative of two independent experiments. Wilcoxon matched-pairs test in the GraphPad Prism 7 software was used to analyze data. The significance was determined by the p-value at the level of 0.05.

**Figure 5.**
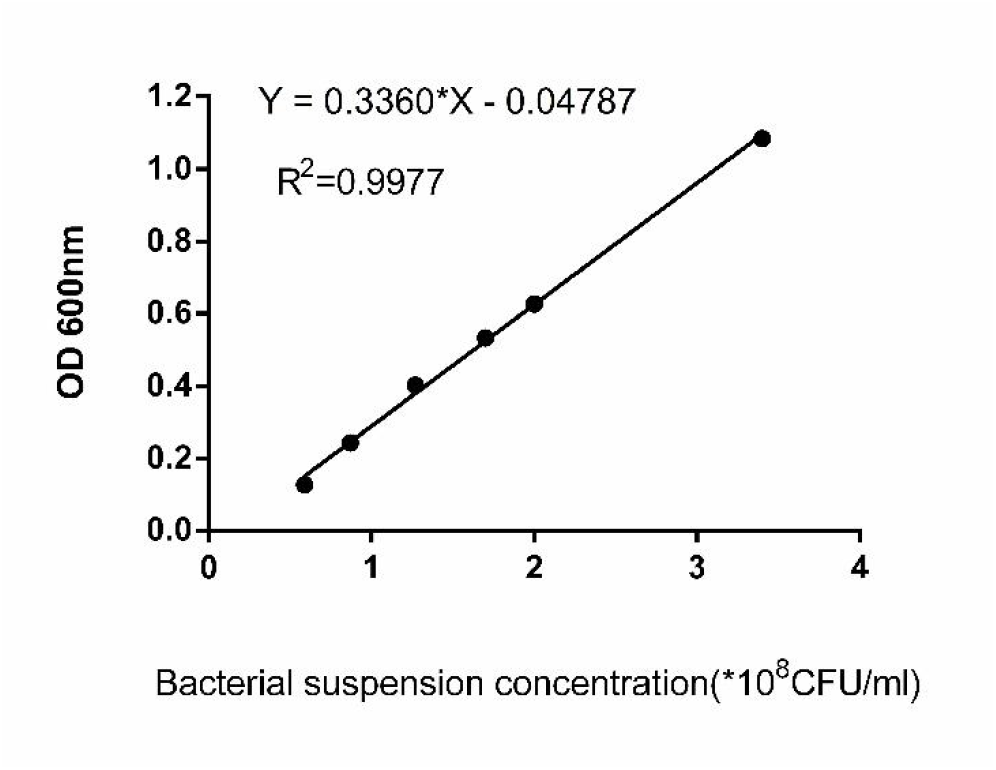
The correlation of the concentration of bacterial suspension and optical density at the wavelength of 600nm. The equation and R-square were calculated by linear regression.

## Discussion

The higher incidence of skin and soft tissue infections (SSTIs) and the emergence of antibiotic-resistant bacteria require the development of novel alternatives against skin bacterial infections (6). Naturally derived antimicrobial peptides, such as caerin peptides, are potential candidates to overcome this challenge since they possess effective antibacterial activity against a broad spectrum of bacterial infections (7). Our previous studies report that caerin 1.9 and its combination with caerin 1.1 have the potential in cancer therapy (18, 23). This study provides additional assessment for the bactericidal properties of caerin 1.9 combined with caerin 1.1 and its potential therapeutic effect on SSTIs. Our results suggested that caerin 1.9 peptide exhibits effective antibacterial activity without inducing resistance, and caerin peptides in temperature sensitive gel formation showed inhibitory effects against bacteria growth in the skin of MRSA-infected mice.

The antimicrobial activity of caerin 1.9 was tested against a range of Gram-positive and Gram-negative bacteria strains, including *S. aureus, P. aeruginosa*, MRSA, *A. Baumannii*, and *S. hemolyiicus*. The MIC values against various bacteria ranged from 3.75 to 15µg/ml (except for *P. aeruginosa*), and it is able to postpone bacterial development at lower concentration (Tabel 1, Figure 1). Moreover, the disc diffusion method was used to compare the susceptibility of caerin peptides with polymyxin B against different bacteria strains. Currently, polymyxin B re-emerged for the treatment of skin-bacterial infections because of the high incidence of multi-drug resistant bacteria and lack of new anti-bacterial agents (24). However, it has limitation on treating certain Gram-positive bacteria strains test in the current study, which required 120µg or above to induce a visible bacterial growth inhibition zone on the agar plates. Our result showed that both caerin 1.1 and caerin 1.9 have an edge over polymyxin B in inhibiting standard *S. aureus*, MRSA, *A. baumannii*, and *S. hemolyiicus* (Table 3).

Caerin 1.1, one representative in the family of caerin peptides, had been demonstrated its antimicrobial activity in multiple studies (12, 14, 25). We found that the combination of caerin 1.1 and caerin 1.9 exhibited an additive effect in inhibiting MRSA and *Acinetobacter baumannii* bacteria growth. Moreover, caerin 1.1 and 1.9 in gel matrix were able to inhibit MRSA growth *in vitro* starting from the concentration of 1.5625mg/ml and displayed a dose-dependent mode up to 12.5mg/ml where the bacteria were completely inhibited (**Figure 2**).

The superficial skin infection model was used to investigate the anti-bacterial activity of caerin gel *in vivo*. This model simulates the situations of SSTI in humans without disrupting the deeper layers of skin (22). It was convenient for evaluating externally-applied antimicrobial agents as dermal antibiotic treatment. Mice presented reduced genetic variations that might influence the results, therefore maximized the reproducibility of the experiments (22, 26). The absorption of caerin gel depends on transdermal delivery. Transdermal delivery of therapeutic peptides permits the direct application of the local infections and increase bioavailability by avoiding hepatic first-pass metabolism and GI absorption (27). It reduces the dosing frequency, hence, lower cost and reduce adverse effects, providing patients with convenient and economical pain-free treatment.

Throughout the 3-day application of caerin gel on the damaged areas, caerin gel significantly inhibited the growth of MRSA in both Balb/c and C57BL/6 mice (Figure 4), and a more severe skin damge model (Figure 3). The gel matrix, poloxamers, are commonly used surfactants in pharmaceuticals, which played a role in stabilizing or dispersing therapeutic agents to enhance dermal absorption. Thus, the poloxamer gel played a role in inhibiting *in vivo* MRSA growth by improving transdermal delivery in this study. The stability of the caerin gel was assessed by Ma et al. in our previous study (18). Although interindividual variations of sensitivity to the caerin therapy existed in the mice, the trend of reduced bacteria growth was observed. However, within the 3-day duration, it was not able to completely eradicate MRSA. Generally, the recovery of skin infection required a longer period, for example, the duration of treatment for impetigo lasted 7 to 14 days depending on severities (28). We are currently investigating the anti-bacterial ativity of the caerin gel at lower concentration and whether the caerin gel can completely clear the skin infection if applied to the skin infection site for a longer period.

Although the mice model resembled human impetigo, it remained some problems that impede the direct application of the results to clinical practice. First of all, these mice were infected with single bacteria, MRSA, but generally, patients in community or hospital settings might be infected with multiple bacteria at the same time (1). Besides, the rate of elimination of microorganisms was faster in animals than in humans due to human complex physiological conditions (26). Therefore, our current results should be translated to realistic clinical trial in future study.

The majority of caerin peptides display effective bactericidal activity through the complementary mechanism of actions; therefore, they are capable to evade the development of resistance (29). Caerin peptides, including caerin 1.9, have similar primary structures based on that of caerin 1.1. Steinborner’s team highlighted that peptides in the caerin 1 family inhibit bacteria cell function with a similar mechanism but various in selectivity towards different bacteria strains (15). The key action for bacterial cell disruption is dependent on membrane interaction, causing pore formation and membrane permeabilization (8, 30). The primary structure (more than 20 amino acids with two Proline residues) of a caerin 1 peptide allows helical structure formation (14, 31). The α-helices aid the peptide insert to lipid membrane for transmembrane pores formation through either barrel-stave mode or toroidal-pore mode (20, 29, 32). Membrane permeabilization leads to the leakage of cell contents, imbalance of homeostasis, membrane dysfunction, and ultimately rapid lysis of bacterial cells. Caerin peptides exhibit alternative mechanism depending on the peptide sequence as well as the concentration of peptide. At high concentrations, caerin 1 peptides may interact with lipid membrane by carpet mechanism, which is more frequent in peptides with less than 20 amino acid residues (7, 32). The affinity and selectivity of peptide-membrane interactions are dependent on the compositions of the lipid membrane and concentration of peptides (30, 33). Therefore, the combined use of caerin 1.1 and caerin 1.9 may increase treatment efficacy and boarder the range of target bacteria.

As a positively charged peptide, caerin peptides can interact with bacterial membranes rather than normal mammalian cell membrane with special selectivity (29). Due to the different compositions and orientations of the lipid membrane, bacterial cells induce stronger negative membrane potential than the eukaryotic cells for peptide selectivity through electrostatic interaction, allowing caerin peptides to bind the bacterial cell membrane in preference (34, 35). Also, high cholesterol contained in mammalian cell membrane aids in maintain membrane stability, which is poor in bacterial cell membrane (29).

In this study, we took an advance to investigate the potential dermal application of caerin 1.1 and caerin 1.9 as a therapy of SSTIs. Nonetheless, some results must be interpreted with caution, and several limitations should be concerned. Due to the restrictions of the MIC experiment, the tested MIC values might vary in every performance. Furthermore, neither the disc diffusion method nor the *in vivo* experiment demonstrates the dose-effect relationship of the antibacterial activity of caerin peptides against MRSA. We are currently using the tape-stripped infected model to determine the minimal dose of caerin peptide gel that inhibit MRSA growth *in vivo* and compare the bacteriacdal efficacy with other antibiotics such as mupiroxin and fusidic acid. Also, the potential toxicity and mechanism of action of the caerin peptides remained undissolved in this study. However, this caerin gel, following further optimization, may provide an alternative method for the management of bacterial infection, especially for multi-drug resistant strains.

In conclusion, caerin 1.9 with a combination of caerin 1.1 have the potential to become a novel antimicrobial agent because of the broad-spectrum antimicrobial activity and limited emergence of bacterial resistance. The peptides in gel formation presented the therapeutic effect of inhibiting bacteria growth in animal models that stimulated human skin bacterial infections. This caerin gel, following further optimization, may provide an alternative method for dermal treatment on skin and soft tissue infections.

## Materials and Methods

### Mice

Six to eight weeks old, specific pathogen**-**free (SPF) adult female C57BL/6 (H-2b) mice and Balb/c mice were ordered from the Animal Resource Centre of Guangdong Province and kept at the Animal Facility of the Foshan First People’s Hospital, Foshan Guangdong, China. Experiments were approved and then performed in compliance with the guidelines of Guangdong Animal Experimentation Ethics Committee (Ethics Approval Number: C202104-1). All mice were kept at clean condition on a 12-hr light/12-hr dark cycle at 22°C and the humidity was 75%. Mice were provided with sterilised standard mouse food and water. Mice were given 1% sodium pentobarbital by *i*.*p*. injection when treatment was performed. At the end of each experiment, mice were sacrificed by CO2 inhalation and confirmed by ceasing breath and heartbeat.

### Caerin peptide and caerin gel preparation

Caerin 1.1 (F1) and Caerin 1.9 (F3) peptides were derived from the Australian tree frog *Litoria splendida*. Caerin 1.1 (GLLSVLGSVAKHVLPHVVPVIAEHL-NH2), caerin 1.9 (GLFGVLGSIAKHVLPHVVPVIAEKL-NH2), and a control peptide which does not have cytotoxic property against a variety of cancerous cells (GTELPSPPSVWFEAEFK-OH) (9), were synthesised by Mimotopes Proprietary Limited, Wuxi, China. The purity of the peptides was >95% as determined by reverse-phase HPLC at Mimotopes. The lipopolysaccharide concentration of caerin 1.1, caerin 1.9 and control peptide (P3) was less than 0.44EU/ml as measured by Kinetic Turbidimetric Assay by Xiamen Bioendo Technology Co., Ltd.

Poloxamer 407 (molecular weight 12600, batch number WPAK592B) and poloxamer 188 (molecular weight 8400, batch number WPAK539B) were purchased from Badische Anilin-und-Soda-Fabrik (BASF; Ludwigshafen, Germany).

The gel matrix was prepared by mixing 46 g of poloxamer 407 and 10 g of poloxamer 188 with 200 ml of distilled water. The preparation was stirred until a white condensation gum matrix was formed and stored at 4°C until the poloxamers were completely dissolved. The caerin peptide gel was prepared by mixing caerin 1.1 and 1.9 with the gel matrix. After the peptides were completely dissolved, the solution was filtered through a 0.22-μm microporous membrane filter to prepare a caerin gel. P3 gel was prepared similarly. The caerin gel, and P3 gels were stored at 4°C until use.

### Bacteria

Standard strains of *Staphylococcus aureus* (S. Aureus, GDM1.441), Copper-Green Pseudomonas (*P. aeruginosa*, GDM1.443), Methicillin-resistant *Staphylococcus aureus* (MRSA, GDM1.1263), Baumann (*Acinetobacter Baumannii*, GDM1.609), and *Streptococcus hemolyiicus* (*S. hemolyiicus*, GDM1.245) were purchased from the Guangdong Microbial Species Conservation Center, Guangdong, China.

Clinical strains of methicillin-resistant *Staphylococcus aureus* (MRSA) were isolated and identified from patients’ specimens of Foshan First People’s Hospital (**Table SX**) by the Department of Pathology of the hospital.

### Bacterial strain resuscitation and preservation

The frozen-dried purchased bacteria were added into a 5 ml MH broth medium (3.65g of Tryptone purchased from Guangdong Ring Kai Microbiology Technology Co., Ltd., dissolved in 100ml of distilled water, 121°C, 20 min autoclaved) and cultured at 37°C for 24 hours. The bacterial suspensions were then inoculated onto a Nutrition agar plate (LS0309, Guangzhou Dijing Microbiology Technology Co., Ltd.) 37°C overnight. The bacterial colonies were picked up using a sterile filter paper and stored at -80°C.

### Establishing the standard curve of bacterial concentration and light optical density

MRSA (GDM1.1263) in the logarithmic growth stage were centrifugated at 8000 rpm for 2 min, washed with phosphate-buffered saline, then resuspended with MH media and inoculated onto a nutrition agar plate (LS0309, Guangzhou Dijing Microbiology Technology Co., Ltd.) to determine the bacteria concentration by counting the colonies. The MRSA suspension was serially diluted and the absorbance was measured by a UV spectrophotometer (UV-7504, Xin Mao, Shanghai, China) at a wavelength of 600 nm. The correlation of bacterial concentration and the value of photo-density was established using linear regression method (Figure S).

### Minimum bacteriological concentration (MIC) of caerin 1.9

The minimum bacteria concentrations (MIC) of caerin 1.9 were measured by using a micro-broth dilution method developed by the Clinical and Laboratory Standards Institute (CLSI) (36). The bacterial suspension with absorbance ranged from 0.08 to 0.10 was prepared and 100 µl of the bacterial suspension to a 96 U-shaped well culture plate, followed by adding 100 µl of different concentrations of caerin 1.9. The final concentrations of the caerin 1.9 were 120, 60, 30, 15, 7.5, 3.75, and 1.875µg/ml respectively. Each concentration of caerin 1.9 was added in triplicate. The equivalent volume of PBS was added as a growth control. The bacteria were cultured at 37°C for 24 hours. The Minimum Inhibitory Concentration (MIC) is defined as the concentration of peptide where bacterial growth is completely inhibited.

### Dynamic bacterial inhibition assay

100μl of bacterial suspensions at the OD value of 0.08-0.1 were added to a 96 well U-shaped plates, followed by adding the same amount of caerin 1.9 solution in triplicate. The final caerin 1.9 concentration is either MIC value or one-fourth of MIC value. PBS was added as a bacterial growth control group. The bacteria were cultured at 37°C for 48 hours and the concentrations were measured at multiple time points by the ELISA plate reader (Multiskan Go, Thermo Scientific, Waltham, MA, USA) at the OD of 600nm.

### Drug resistance induction

*P. aeruginosa* and MRSA were prepared with dilution, and then cultured in a 96-well U shape plate containing 1/4MIC concentration of caerin 1.9, Tazocin, or PBS. Every three days, the cultured bacteria were transferred to new media containing the same amount of caerin 1.9 or Tarocin. After 30 rounds of culture, the MIC of caerin 1.9 or Tazocin on the two bacteria was measured as described above. The MIC values before and after the treatment were compared.

### Antimicrobial susceptibility test of caerin 1.1 and caerin 1.9

The antimicrobial activity of caerin 1.1 and caerin 1.9 were evaluated using the disc diffusion method according to the guidelines of the Clinical and Laboratory Standards Institution (CLSI) (37). Firstly, the nutrition agar plates were inoculated with standardized bacteria strains. Then, 6mm filter paper discs which were carpeted with 120µg of caerin 1.1, caerin 1.9, or polymyxin B were placed on the agar surface. The plates were cultured at 37°C for 24 hours to allow the formation of the inhibition growth zone. The inhibition zone diameter was measured using a vernier caliper.

### Caerin 1.1 and caerin 1.9 combination against MRSA and Baumannii

The combination of caerin 1.1 and caerin 1.9 against MRSA and *P. Baumannii* were tested with a microdilution checkerboard assay described elsewhere (21). Firstly, the serially diluted bacterial suspensions were cultured in a 96 well U-shaped culture plate in the presence of caerin 1.1 and 1.9. The final concentrations of caerin 1.1 and 1.9 in the plate were from 2×MIC to l/64 MIC. For wells along the x-axis, caerin 1.9 was added, while caerin 1.1 was added along the y-axis. Each well within this 8×8 checkerboard contained a combination of caerin 1.1 and 1.9 at different concentrations. The fractional inhibitory concentration index (FICI) for a well was calculated as follows,

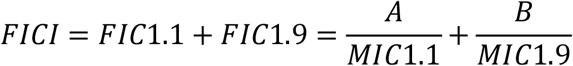

Where, A is the concentration of caerin 1.1 in a given well along the inhibition-no-inhibition interface; MIC1.1 is the MIC of caerin 1.1 alone; FIC1.1 is the fractional inhibitory concentration of caerin 1.1; and B, MIC1.9, and FIC1.9 are defined in the same fashion for caerin 1.1. FICI ≤1.0 implies additive effect or synergy.

### Skin infection model

Six- to eight-week-old female Balb/c mice with their fur on the back removed were anesthetized and the skin was cut into an oval shape of around 1×2cm. A bacterial infection was initiated by placing MRSA suspension (containing 5×10^6^ CFU) onto each damaged area, followed by application of 20µl of caerin gel to the left side of the striped area and 20µl of poloxamer gel to the right side. The treatment was performed twice at 4 hour and 16 hour respectively. At 24 hour, pus and secretions on each damaged area were collected and diluted serially with normal saline, and then inoculated onto a nutrition agar plate for culture overnight. The colonies were counted and compared.

### Tape-stripping infection model

A tape-stripping infection model was used to investigate the antibacterial abiity of caerin 1.1 and 1.9 gel (22). Briefly, six-to eight-week-old female Balb/c mice or C57BL/6 mice were anesthetized. After the removal of the fur on the back, the skin was stripped with an elastic bandage (Smith & Nephew Medical, Hull, United Kingdom). Two areas of 1cm×2cm on both left and right sides of the torso was tape stripped. After stripping, 10µl of MRSA suspension (GIM1.1263, 5×10^6^ CFU/ml) was applied for each stripped surface. Four hours after the skin infection, 20µl of caerin 1.1 and 1.9 gel containing 12.5mg/ml each was applied to the left side of the striped area, while the same amount of poloxamer gel to the right side. In another group, 20µl of P3 gel of 12.5mg/ml was applied to the left, and poloxamer gel to the right. Mice in the infection control group were given 20µl of normal saline. Each treatment was performed twice daily for consecutive three days with a total of five times. On the fourth day, the striped areas were removed and subsequently homogenized by glass homogenizer and diluted with normal saline, followed by inoculation on nutrition agar plates (LS0309, Guangzhou Dijing Microbiology Technology Co., Ltd.) for culture for 24 hours before the colonies were counted.

### Statistical analysis

Paired student t-test statistical analysis was performed to evaluate the in vivo bacteria growth inhibition using GraphPad Prism 7 software. All experimental data were analyzed and graphs were plotted in the same software. The significant means were determined at the probability level of 0.05.

## Acknowlegdement

This study was partly supported by the National Natural Science Foundation of China (31971355), and the DengFeng project of Foshan First People’s Hospital (2019A008). The funders had no role in study design, data collection and interpretation, or the submission of the work for publication. All authors read and agreed to the published version of the manuscript.

